# The Positive End of the Polygenic Score Distribution for ADHD: A Low Risk or a Protective Factor?

**DOI:** 10.1101/611897

**Authors:** James J. Li

## Abstract

**Background:** Polygenic scores (PGS) are widely used to characterize genetic liability for heritable mental disorders, including attention-deficit/hyperactivity disorder (ADHD). However, little is known about the effects of a low burden of genetic liability for ADHD, including whether this functions as a low risk or protective factor for ADHD and related functional outcomes in later life. The current study examines the association of low ADHD PGS and functional outcomes in adulthood.

**Methods:** Participants were from Wave IV of the National Longitudinal Study of Adolescent to Adult Health (Add Health) (*N*=7,088; mean age=29, *s.d*.=1.74). ADHD PGS was computed from an existing genome-wide association study, and adult functional outcomes, including cognition, educational attainment, mental health and physical health were assessed during in-home interviews.

**Results:** Individuals at the lowest end of the ADHD PGS distribution (i.e., lowest 20^th^ percentile) had the lowest probabilities of ADHD, exhibiting a 17-19% reduction in risk for ADHD relative to the observed 8.3% prevalence rate of ADHD in Add Health. Furthermore, individuals with low ADHD PGS had higher cognitive performance, greater levels of educational attainment, and lower BMI relative to individuals representing the rest of the ADHD PGS distribution, including those who were in the medium and high PGS groups.

**Conclusions:** Findings indicate that psychiatric PGS likely capture far more than just the risk and the absence of risk for a psychiatric outcome; where one lies along the PGS distribution may predict diverging functional consequences, for better and for worse.

---

Attention-deficit/hyperactivity disorder (ADHD) is a neurodevelopmental disorder with an estimated prevalence of 8.9% among children (aged 6 to 11) in the United States (Danielson et al., 2018). It is associated with a range of negative functional outcomes in adulthood, including poor cognitive functioning (Boonstra et al., 2005), low educational attainment (Kuriyan et al., 2013), depression and substance misuse (Agnew-Blais et al., 2018; Lee et al., 2011), involvement in the criminal justice system (Fletcher and Wolfe, 2009), increased stress (Combs et al., 2015) and poor physical health (Brook et al., 2013; Kuriyan et al., 2013). Etiologically, ADHD is among the most heritable of the psychiatric disorders; twin studies have estimated that between 70-75% of the variance in ADHD dimensions can be accounted for by genetic differences in the population (Chang et al., 2013; Faraone and Larsson, 2018; Nikolas and Burt, 2010). Recently, this heritability has been shown to be highly polygenic, reflecting many genes of individually small effects (Demontis et al., 2019; Franke et al., 2009; Larsson et al., 2014; Zayats et al., 2015). As such, studies are increasingly using polygenic scores (PGS) (Dudbridge, 2013) to characterize the aggregate genetic liability for ADHD.

PGS are typically computed as the linear composite of alleles one carries across single nucleotide polymorphisms (SNPs) tested in association with a psychiatric trait, with each SNP weighted according to genome-wide association study (GWAS) summary statistics on the trait in question (Anderson et al., 2019). One consistent observation of psychiatric PGS is that they are distributed normally in population-based samples (Krapohl et al., 2016), reflecting the continuum of genetic risk for psychopathology in a population (Plomin et al., 2009). Whereas individuals on the right-tail (e.g., >20^th^ percentile) are often characterized as “high risk” (Khera et al., 2018; Torkamani et al., 2018), individuals at the left-tail (e.g., <20^th^ percentile) are traditionally considered a “low risk” group (Torkamani et al., 2018). However, the term “low risk” for this subgroup is ambiguous and questionable, especially given that few studies have empirically examined the implications of having low PGS. For instance, low risk could equate to the baseline of risk for the disorder observed in the population. Low risk could also indicate a *reduction* of risk for the disorder relative to the population baseline, perhaps reflecting a biologically protective effect of having few risk alleles. In general, few researchers have been “thinking positively” (pp. 875, Plomin et al., 2009) about PGS, and thus, may have been neglecting potentially important functional consequences of being in the low end of the PGS distribution.

Studies have reliably shown that high PGS is associated with greater risk for ADHD and related negative outcomes (Demontis et al., 2019; Du Rietz et al., 2017; Groen-Blokhuis et al., 2014; Hamshere et al., 2013; Riglin et al., 2016; Stergiakouli et al., 2015). Conversely, low PGS has also been shown to confer reduced risk for ADHD relative to the higher PGS groups (Demontis et al., 2019; Wray et al., 2018). However, it is questionable whether low PGS also confers a *reduction* in risk for ADHD relative to individuals in the general population, such as those with medium PGS. In a study of 1,985 depressed patients and controls from the Netherlands, the odds of having a major depressive disorder (MDD) was highest among those who had the highest MDD PGS and experienced severe childhood maltreatment (relative to individuals experiencing moderate or no/low maltreatment) (Peyrot et al., 2014). However, individuals at the lowest end of the MDD PGS distribution showed low odds of MDD regardless of maltreatment severity (Peyrot et al., 2014). Notably, this finding was inconsistent with findings from a similar study featuring depressed patients and controls from the United Kingdom (Mullins et al., 2016).

Krapohl and colleagues (2016) showed that individuals in the lowest septile PGS for educational attainment had the greatest amount of parent-reported behavior problems relative to individuals in the highest PGS septile, who had the fewest behavior problems (Krapohl et al., 2016). However, no associations were detected between low (or high) psychiatric PGS, including for ADHD, with various behavioral and socioemotional outcomes. Notably, this study was limited by the underpowered GWAS for psychiatric outcomes relative to the educational attainment GWAS. Given that a significantly higher-powered GWAS is now available for ADHD, a re-examination of this association is warranted.

Despite the ascendancy of PGS in genetic association studies, most studies in this emerging literature have ignored the functional implications of having low psychiatric PGS (Anderson et al., 2019; Bogdan et al., 2018). This study used data from a population-based dataset to investigate the association of the ADHD PGS distribution as it pertains to ADHD and functional outcomes associated with ADHD, including educational attainment, cognition, mental and physical health. In line with the hypothesis that low ADHD PGS may confer a protective effect on ADHD and related functional outcomes, individuals with low ADHD PGS were expected to have a reduced risk of ADHD, relative to 1) those at medium and higher ADHD PGS levels, and 2) the observed population prevalence rate of ADHD in Add Health. Furthermore, individuals with low ADHD PGS were also expected to have superior functional outcomes in adulthood relative to those with medium and high ADHD PGS.

## Methods

### Participants

Data were from the National Longitudinal Study of Adolescent to Adult Health (Add Health), a stratified sample of adolescents in grades 7-12 from high schools across the U.S. Data were collected from adolescents, parents, fellow students, school administrators, siblings, friends and romantic partners across four waves: Wave I (1994-1995, grades 7-12, *N*=20,745), Wave II (1995-1996, grades 8-12, *N*=14,738), Wave III (2001-2002, ages 18-26, *N*=15,197), and Wave IV (2007-2008, ages 24-32, *N*=15,701). Wave IV phenotypic data were used given the focus on adult outcomes as a function of childhood ADHD. The current analyses were performed for individuals where both genotypic and phenotypic information were available (*N*=7,088). Within the genotypic subsample, the mean age at Wave IV was 29.00 (*s.d*.=1.74), 46% of this sample was male, and the racial-ethnic composition was 63.6% Caucasian (including Hispanic), 20.7% African American, .2% Native American, 5.1% Asian, and 10.3% “Other.” Although there were a few statistical differences between the genetic and non-genetic Add Health subsamples across demographic variables (see Online Supplement, eTable 1), these differences were very small in terms of their effect sizes.

### Measures

#### ADHD

Childhood ADHD symptoms were self-reported by the participant retrospectively (when the participants were between ages 5 and 12) at Wave III, keyed to the *Diagnostic and Statistical Manual of Mental Disorders* (American Psychiatric Association, 2013). 17 items were rated on a 4-point Likert scale regarding how often the symptom “best describes your behavior when you were [between 5 and 12].” Each item was dichotomized to indicate the presence (i.e., *often* or *very often* response) or absence (i.e., *never* or *sometimes* response) of the symptom. In line with the DSM, diagnostic criteria for ADHD was defined as having 6 or more symptoms of inattention and/or 6 or more symptoms of hyperactivity/impulsivity.

#### Cognitive Ability

Cognitive ability was assessed via the Add Health Picture Vocabulary Test (AHPVT) at Wave I. The task measures receptive vocabulary, verbal ability and scholastic aptitude. For this test, the interviewer read a word aloud and the participant selected an illustration that best fit its meaning. Each word had four simple, black-and-white illustrations arranged in a multiple-choice format. The standardized score for AHPVT was used.

#### Educational Attainment

Educational Attainment was assessed at Wave IV through the following question: “What is the highest level of education that you have achieved to date?” The scale ranged from 1 (“8^th^ grade or less”) to 10 (“some graduate training beyond a master’s degree”).

#### Mental Health and Behavior

All mental health and behavior outcomes were assessed at Wave IV. Depression symptoms were measured using an abbreviated version of the Center for Epidemiologic Studies Depression Scale (CES-D) (Radloff, 1977). Lifetime DSM-IV criteria for alcohol abuse or dependence was assessed as the presence of at least 1 of the 4 items pertaining to alcohol abuse, and/or 3 of the 7 items pertaining to alcohol dependence occurring together in a 12-month period. Lifetime DSM-IV criteria for “other drug” abuse and dependence were assessed using the same criteria, but for illicit substances. ‘Ever arrested’ was measured by whether the participant responded affirmatively to the question: “have you ever been arrested” and/or whether the Wave IV interview was conducted with the participant while in prison. Perceived stress was measured via an abbreviated 4-item version of the Cohen’s Perceived Stress Scale (Cohen et al., 1983), which assessed perception of stress across various life contexts. Items were rated on a 5-point Likert scale, where 0=“never” and 4=“very often.”

#### Physical Health

All physical health outcomes were assessed at Wave IV. Body mass index (BMI) classification was measured on a 1-6 scale, where 1=underweight (BMI<18.5), 3=overweight (BMI=20-30), and 6=obese III (BMI>40). Stage 2 hypertension was present if the participant responded affirmatively to the question: “has a doctor, nurse or other health care provider ever told you that you have or had: high blood pressure or hypertension, systolic blood pressure, and diastolic blood pressure?” High blood cholesterol was present if the participant responded affirmatively to the question: “has a doctor, nurse or other health care provider ever told you that you have or had: high blood cholesterol or triglycerides or lipids?”

#### Genotyping and Quality Control

Saliva were obtained from participants at Wave IV. Genotyping was done on the Omni1-Quad BeadChip and the Omni2.5-Quad BeadChip. Add Health European genetic samples were imputed on Release 1 of the Human Reference Consortium (HRS r1.1). Non-European samples were imputed using the 1000 Genomes Phase 3 reference panel. Of 606,673 variants, 13,721 were removed with a per-variant missing call rate filter of 0.02; 245,589 were removed with a Hardy-Weinberg Equilibrium filter of 0.0001, and 609 were removed with a minor allele frequency filter of 0.01, leaving 346,754 SNPs carried through to imputation. Additional details of the quality control are available online (https://www.cpc.unc.edu/projects/addhealth/documentation/guides).

#### Polygenic Scores (PGS)

PGS were computed as the linear composite of SNPs associated with ADHD, weighted by each SNPs effect size according to meta-analytic GWAS (Demontis et al., 2019). This GWAS featured 55,374 individuals (20,183 cases and 35,191 controls) from 12 studies of mixed (but predominantly European) ancestries and a replication sample of 93,916 individuals from two mixed ancestry cohorts. This study used an GWAS p-value threshold of *p*=1 rather than an empirical *p*-value threshold to minimize potential bias due to overfitting (Benjamini et al., 2001). PGS were then standardized according to genetic ancestry groups in Add Health (Braudt and Harris, 2018). Additional controls for the population stratification were done by covarying the first 10 ancestry-specific principal components (PC) of the genetic data in the analyses (Price et al., 2006).

#### Statistical Analysis

First, a logistic regression was modeled where ADHD PGS (measured continuously and categorically) was regressed on ADHD diagnostic status, controlling for age, biological sex and genetic PCs. Predicted and observed probabilities of ADHD were computed for each category of the PGS (i.e., binned into 20^th^ percentiles; see Figure 1). Then, low (<20^th^ percentile), medium (21^st^ – 70^th^ percentiles) and high (>80^th^ percentile) PGS groups were compared across adult functional outcomes. These group comparisons were conducted using parametric and nonparametric tests: a one-way ANOVA was performed to examine differences on the AHPVT standardized scores (a normally distributed quantitative outcome), Kruskal-Wallis H tests were used to test for differences on ordinal or quantitative discrete/non-normal outcomes (i.e., educational attainment, depression symptoms, perceived stress, and BMI) and chi-square tests were used for binary outcomes (e.g., DSM-IV alcohol abuse/dependence, illicit drug abuse/dependence, ever arrested, hypertension and high blood cholesterol). Pairwise contrasts were probed between each PGS group on the outcome variables. The alphas were set to .005 to correct for multiple testing (i.e., .05 divided by 11 dependent variables). Bivariate correlations between PGS and ADHD as well as all other dependent variables are in eTable 2. Due to the stratified and clustered nature of the sampling performed in Add Health, sampling weights were applied in all analyses.

**Figure 1.**
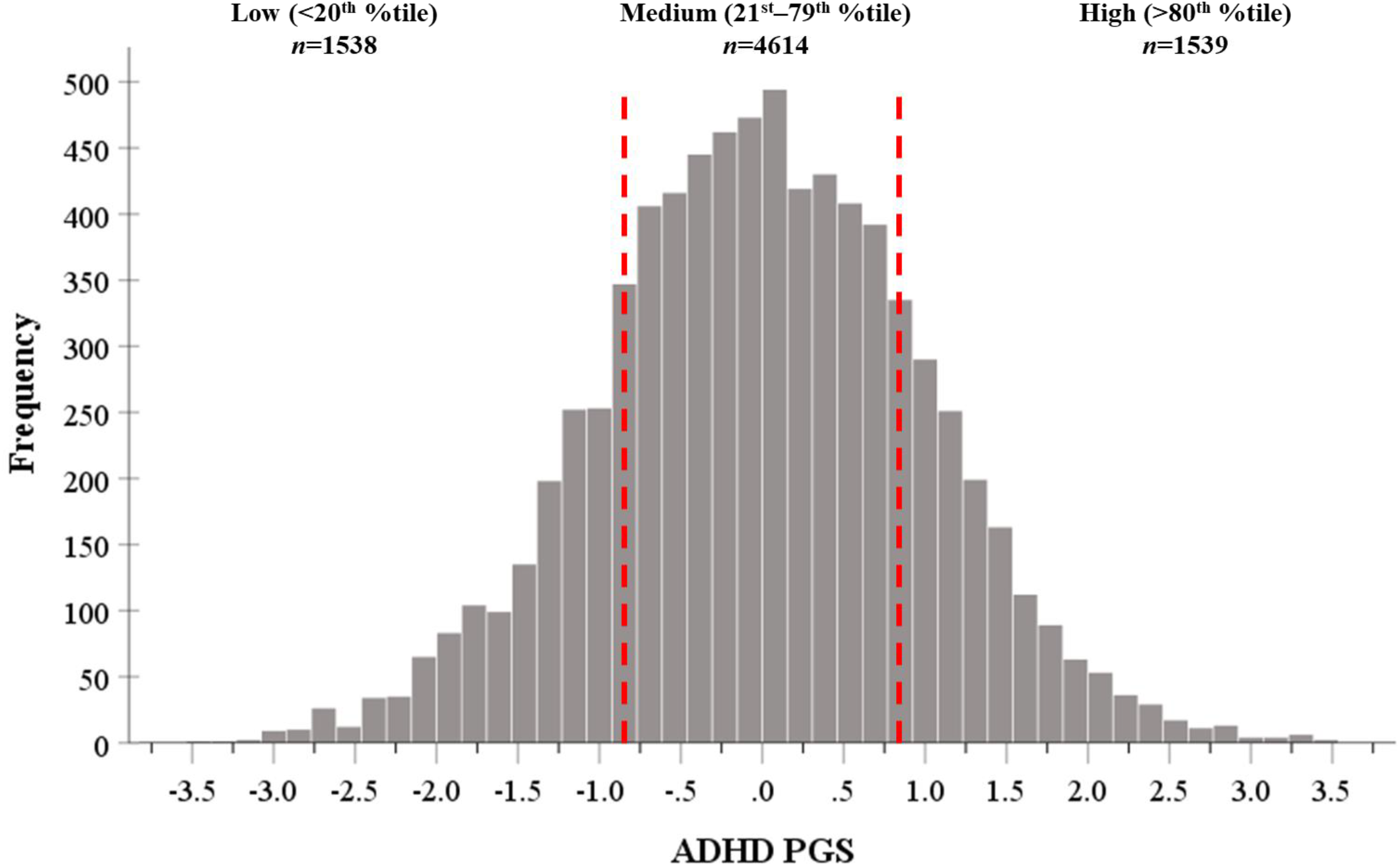
The ADHD PGS distribution in Add Health. Red lines represent thresholds at <20^th^ percentile (ADHD PGS ≤ −.83) and >80^th^ percentile (ADHD ≥ .85) of the ADHD PGS distribution, respectively.

## Results

### PGS and ADHD Diagnostic Status

As expected, ADHD PGS was positively associated with ADHD diagnostic status (OR=1.22, *p*<.001, 95% CI=1.10-1.36). Figure 1 shows the predicted and observed probabilities of ADHD by ADHD PGS group, ranging the bottom 20^th^ percentile (PGS≤-.83) of PGS to the top 20^th^ percentile (PGS≥.85). ADHD probabilities significantly differed by PGS group (χ^2^(2)=22.21, *p*<.001). Individuals in the lowest 20^th^ percentile of ADHD PGS (i.e., low PGS) had lower probabilities of ADHD than those in the middle percentiles (i.e., medium PGS; χ^2^(1)=6.04, *p*=.01) and those in the highest 20^th^ percentile (i.e., high PGS; χ^2^(4)=20.24, *p*<.001), although it should be noted that the pairwise contrast between the low and medium PGS did not reach significance with a Bonferroni correction (*p*<.005). Furthermore, those in the medium PGS group had lower probabilities of ADHD than those in the high PGS group (χ^2^(1)=9.69, *p*=.002).

For those in the low PGS group, the predicted probability of ADHD was 6.77% (*s.e*.=.62, 95% CI=5.54-7.99%) and the observed probability was 6.87% (*s.e*.=.83, 95% CI=5.39-8.71%). Overall, individuals in the low PGS group had a roughly 17-19% reduction in risk for ADHD relative to the Add Health ADHD prevalence rate of 8.3%.

Individuals in the medium PGS group had a predicted ADHD probability of 8.78% (*s.e*.=.49, 95% CI=7.82-9.75%) and an observed ADHD probability of 8.78% (*s.e*.=.65, 95% CI=7.58-10.14%). As expected, those in the medium PGS group had an ADHD risk profile that was approximately equivalent to the general population as well as the Add Health ADHD prevalence rate.

Individuals in the high PGS group had the highest probabilities of ADHD; the predicted probability of ADHD was 11.22% (*s.e*.=.87., 95% CI=9.49-12.95%) and the observed probability was 11.14% (*s.e*.=1.10, 95% CI=9.14-13.53%). Being in the top 80^th^ percentile of polygenic risk for ADHD conferred a 35% increase in risk for ADHD relative to the Add Health ADHD prevalence rate of 8.3%.

### PGS Groups and Adult Functional Outcomes

Parametric and nonparametric tests were performed to compare PGS group differences across cognition, educational attainment, mental health and behavior, and physical health outcomes (Table 1 and Figure 2). There were significant group differences for most of the outcomes examined, with the exception of DSM-IV alcohol abuse/dependence (χ^2^(2)=2.08, *p*=.350), hypertension (χ^2^(2)=5.72, *p*=.060), and high blood cholesterol (χ^2^(2)=2.85, *p*=.240).

**Table 1.** Tests comparing PGS status across functional outcomes

| Dependent Variable (range) | PGS Status | Mean | s.e. | Test Statistic |  |  | 95% Confidence Interval |  |
| --- | --- | --- | --- | --- | --- | --- | --- | --- |
| | | | | $F, \chi^2$ or $H$ | $p^*$ | $df$ | Lower | Upper |
| Cognition |  |  |  |  |  |  |  |  |
| AHPVT standardized score (50-141) | Low | 104.13 | .72 | $F=24.56$ | <.001 | 2 | 102.70 | 105.55 |
|  | Medium | 101.40 | .55 |  |  |  | 100.32 | 102.49 |
|  | High | 99.06 | .64 |  |  |  | 97.80 | 100.32 |
| Educational Attainment |  |  |  |  |  |  |  |  |
| Highest degree attained (1-10) | Low | 5.87 | .11 | $H=109.30$ | <.001 | 2 | 5.66 | 6.08 |
|  | Medium | 5.54 | .07 |  |  |  | 5.40 | 5.70 |
|  | High | 5.09 | .10 |  |  |  | 4.89 | 5.29 |
| Mental Health and Behavior |  |  |  |  |  |  |  |  |
| CES-D depression symptoms (0-15) | Low | 6.17 | .16 | $H=11.71$ | <.001 | 2 | 5.85 | 6.49 |
|  | Medium | 6.73 | .11 |  |  |  | 6.52 | 6.93 |
|  | High | 7.00 | .21 |  |  |  | 6.59 | 7.41 |
| DSM-IV alcohol abuse/dependence (0-1) | Low | .32 | .02 | $\chi^2=2.08$ | .350 | 2 | .29 | .35 |
|  | Medium | .29 | .01 |  |  |  | .27 | .31 |
|  | High | .25 | .02 |  |  |  | .21 | .29 |
| DSM-IV illicit drug abuse/dependence (0-1) | Low | .08 | .01 | $\chi^2=13.12$ | <.001 | 2 | .06 | .10 |
|  | Medium | .08 | .00 |  |  |  | .07 | .09 |
|  | High | .10 | .01 |  |  |  | .08 | .12 |
| Ever arrested (0-1) | Low | .26 | .01 | $\chi^2=28.32$ | <.001 | 2 | .24 | .29 |
|  | Medium | .29 | .01 |  |  |  | .27 | .31 |
|  | High | .34 | .02 |  |  |  | .31 | .38 |
| Perceived Stress (0-16) | Low | 4.56 | .11 | $H=13.91$ | <.001 | 2 | 4.34 | 4.77 |
|  | Medium | 4.85 | .08 |  |  |  | 4.70 | 5.00 |
|  | High | 5.02 | .11 |  |  |  | 4.79 | 5.24 |

Table 1. Tests comparing PGS status across functional outcomes (continued from previous page)
|  |  |  |  | Test Statistic |  |  | 95% Confidence Interval |  |
| --- | --- | --- | --- | --- | --- | --- | --- | --- |
| Dependent Variable | PGS Status | Mean | s.e. | *F*, $\chi^2$ or *H* | *p* * | *df* | Lower | Upper |
| Physical Health |  |  |  |  |  |  |  |  |
| BMI (1-6) | Low | 3.11 | .04 | $H=37.70$ | <.001 | 2 | 3.04 | 3.18 |
|  | Medium | 3.33 | .03 |  |  |  | 3.27 | 3.39 |
|  | High | 3.47 | .06 |  |  |  | 3.36 | 3.58 |
| Hypertension stage 2 (0-1) | Low | .12 | .01 | $\chi^2=5.72$ | .060 | 2 | .09 | .14 |
|  | Medium | .14 | .01 |  |  |  | .13 | .16 |
|  | High | .14 | .01 |  |  |  | .12 | .17 |
| High blood cholesterol (0-1) | Low | .08 | .01 | $\chi^2=2.85$ | .240 | 2 | .06 | .10 |
|  | Medium | .07 | .01 |  |  |  | .06 | .08 |
|  | High | .09 | .01 |  |  |  | .07 | .11 |

\*P-values are reported to the thousandsth decimal point due to the Bonferonni correction threshold of $p<.005$ .
Note. Results show parametric and nonparametric tests of PGS group comparisons on functional outcomes. A one-way ANOVA was performed to examine differences on continuous and normally distributed outcomes. Kruskal-Wallis H tests were used to test for differences on ordinal or continuous outcomes, and chi-square tests were used for binary outcomes. “Low” PGS is the lowest quintile (<20<sup>th</sup> percentile) of the PGS distribution, “Medium” PGS is the combination of the second, third, and fourth quintiles (21<sup>st</sup> – 79<sup>th</sup> percentile) of the PGS distribution, “High” PGS is the fifth quintile (>80<sup>th</sup> percentile) of the PGS distribution.

**Figure 2.**
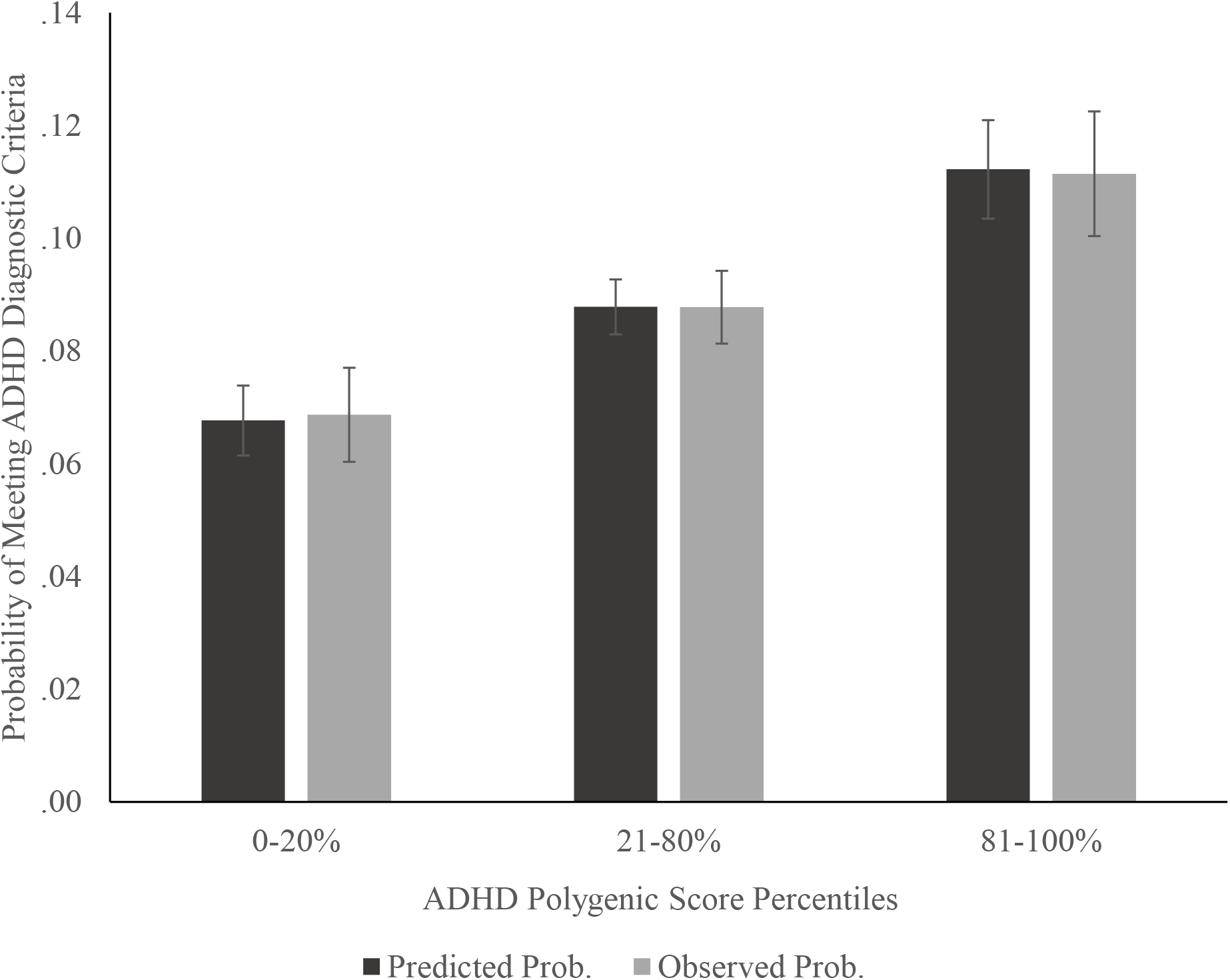
Predicted and observed probabilities of meeting diagnostic criteria for ADHD in Add Health by ADHD PGS. Figure shows the predicted and observed probabilities of meeting diagnostic criteria for ADHD in Add Health by ADHD PGS groups, according to a binary logistic regression that controlled for age, biological sex, and the first 10 principal components of the genetic information (i.e., genetic ancestry). Error bars reflect standard errors.

**Figure 3.**
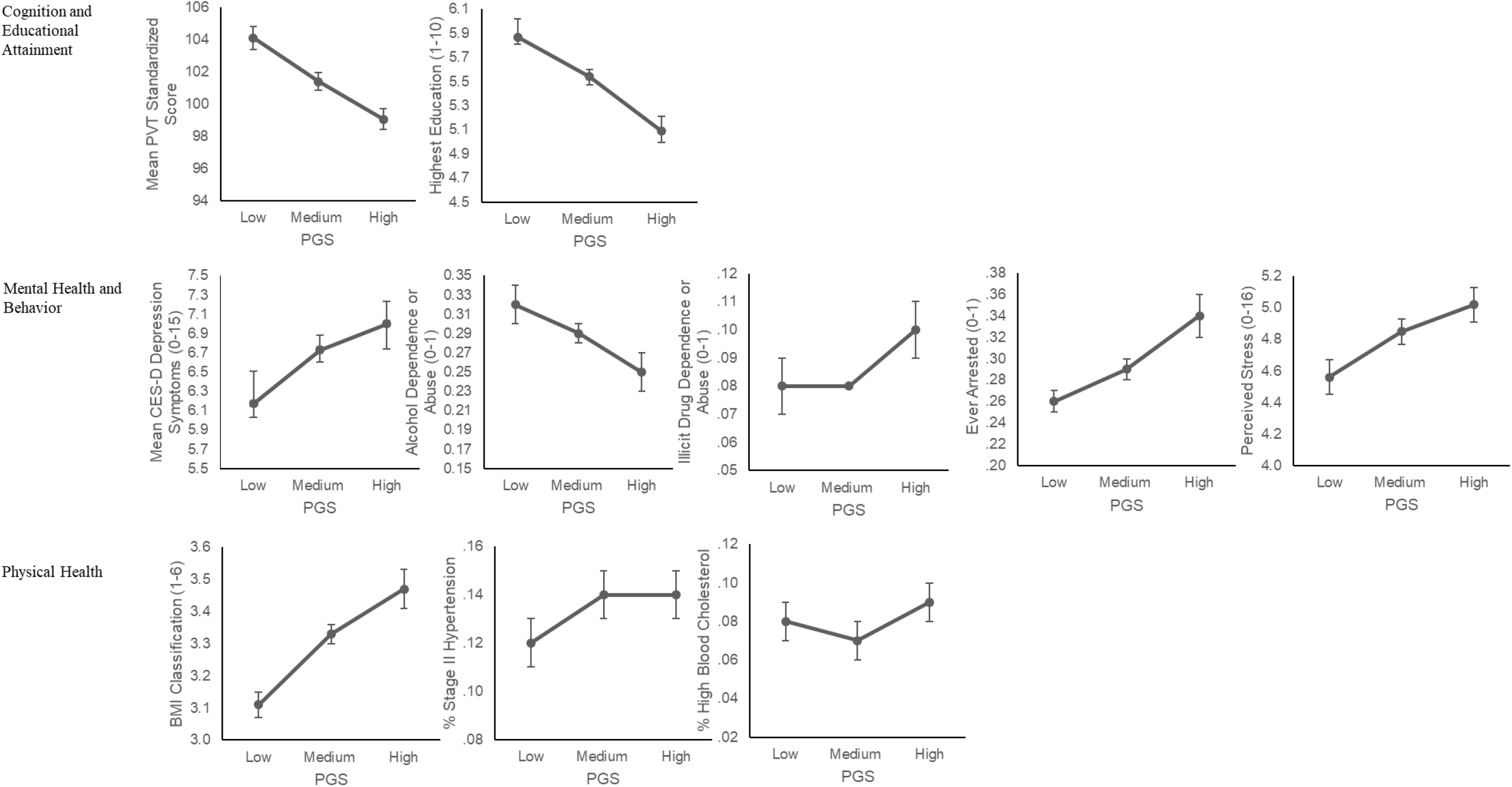
Associations between ADHD PGS levels and functional outcomes. Plots of the observed means for each dependent variable by each ADHD PGS level, obtained from a multivariate analysis of variance controlling for age, biological sex, and the first 10 principle components of the genetic information (i.e., genetic ancestry). “Low” PGS is the lowest quintile (<20^th^ percentile) of the ADHD PGS distribution, “Medium” PGS is the combination of the second, third, and fourth quintiles (21^st^ – 79^th^ percentile) of the ADHD PGS distribution, “High” PGS is the fifth quintile (>80^th^ percentile) of the ADHD PGS distribution. Standard errors are shown on each figure.

Pairwise contrasts (Table 2) showed, not surprisingly, that low and high PGS individuals differed most significantly (after Bonferroni correction) across the functional outcomes, where low PGS individuals had significantly higher AHPVT standardized scores and educational attainment, significantly fewer depression symptoms, and significantly lower rates or levels of illicit drug abuse/dependence, being ever arrested, perceived stress, and BMI relative to their high PGS counterparts. These groups did not differ on their alcohol abuse/dependence rates, hypertension, or on high blood cholesterol.

**Table 2.** Pairwise contrasts of PGS groups on dependent variables

| PGS Group Contrasts<br>(dummy code) | Dependent Variable | Test Statistic |  |  |
| --- | --- | --- | --- | --- |
|  |  | *T, $\chi^2$ or H* | *df* | *p\** |
| Low (0) versus Medium (1) | Cognition |  |  |  |
|  | AHPVT standardized score | *T*=5.52 | 5871 | <.001 |
|  | Educational Attainment |  |  |  |
|  | Highest degree attained | *H*=36.14 | 1 | <.001 |
|  | Mental Health and Behavior |  |  |  |
|  | CES-D depression symptoms | *H*=2.56 | 1 | .109 |
| | DSM-IV alcohol abuse/dependence | $\chi^2$=.81 | 1 | .369 |
| | DSM-IV illicit drug abuse/dependence | $\chi^2$=.24 | 1 | .625 |
| | Ever arrested | $\chi^2$=2.45 | 1 | .118 |
|  | Perceived stress | *H*=2.85 | 1 | .091 |
|  | Physical Health |  |  |  |
|  | BMI | *H*=21.56 | 1 | <.001 |
| | Hypertension stage 2 | $\chi^2$=4.14 | 1 | .042 |
| | High blood cholesterol | $\chi^2$=.48 | 1 | .487 |
| Low (0) versus High (2) | Cognition |  |  |  |
|  | AHPVT standardized score | *T*=6.75 | 2934 | <.001 |
|  | Educational Attainment |  |  |  |
|  | Highest degree attained | *H*=107.24 | 1 | <.001 |
|  | Mental Health and Behavior |  |  |  |
|  | CES-D depression symptoms | *H*=8.97 | 1 | .003 |
| | DSM-IV alcohol abuse/dependence | $\chi^2$=2.08 | 1 | .149 |
| | DSM-IV illicit drug abuse/dependence | $\chi^2$=9.05 | 1 | .003 |
| | Ever arrested | $\chi^2$=24.13 | 1 | <.001 |
|  | Perceived stress | *H*=13.61 | 1 | <.001 |

Table 2. Pairwise contrasts of PGS groups on dependent variables (continued from previous page)
| PGS Group Contrasts | Dependent Variable | Test Statistic |  |  |
| --- | --- | --- | --- | --- |
|  |  | *T, $\chi^2$ or *H** | *df* | *p* * |
| Low (0) versus High (1) | Physical Health |  |  |  |
| | BMI | $H=35.72$ | 1 | <.001 |
| | Hypertension stage 2 | $\chi^2=5.19$ | 1 | .023 |
| | High blood cholesterol | $\chi^2=.60$ | 1 | .438 |
| Medium (1) versus High (2) | Cognition |  |  |  |
| | AHPVT standardized score | $T=2.85$ | 5873 | .004 |
|  | Educational Attainment |  |  |  |
| | Highest degree attained | $H=46.82$ | 1 | <.001 |
|  | Mental Health and Behavior |  |  |  |
| | CES-D depression symptoms | $H=4.23$ | 1 | .040 |
| | DSM-IV alcohol abuse/dependence | $\chi^2=.75$ | 1 | .386 |
| | DSM-IV illicit drug abuse/dependence | $\chi^2=11.00$ | 1 | .001 |
| | Ever arrested | $\chi^2=20.37$ | 1 | <.001 |
| | Perceived stress | $H=7.72$ | 1 | .005 |
|  | Physical Health |  |  |  |
| | BMI | $H=7.81$ | 1 | .005 |
| | Hypertension stage 2 | $\chi^2=.53$ | 1 | .467 |
| | High blood cholesterol | $\chi^2=2.79$ | 1 | .095 |

\*P-values are reported to the thousandsth decimal point due to the Bonferonni correction threshold of $p<.005$ .

Interestingly however, low PGS individuals also differed significantly from medium PGS individuals on several functional outcomes, suggesting the possibility of a protective effect of low PGS. In particular, low PGS individuals had significantly higher AHPVT standardized scores and greater levels of educational attainment, as well as significantly lower levels of BMI. Low PGS individuals did not significantly differ from their medium PGS counterparts on mental health and behavioral outcomes (i.e., depression symptoms, alcohol abuse/dependence, illicit drug abuse/dependence, rates of being ever arrested, perceived stress) and on other indices of physical health (i.e., hypertension and high blood cholesterol).

Finally, and also notably, medium PGS individuals differed from high PGS individuals across all the major functional domains. Medium PGS individuals had higher AHPVT standardized scores and educational attainment, lower rates of illicit drug abuse/dependence, being ever arrested, and perceived stress, and also lower BMI relative to their high PGS counterparts.

### Secondary Analyses

ADHD PGS may not be generalizable to non-European populations given that the GWAS was performed on predominantly European-ancestry populations (Demontis et al., 2019). Although these analyses include safeguards against population stratification (i.e., controlling for 10 PCs) and standardized PGS by genetic ancestry, parallel analyses were conducted for the European-only subsample of Add Health (*n*=4,506) to determine whether the results were only specific to European-ancestry individuals in the sample. ADHD PGS was significantly predictive of ADHD in the European-subsample (OR=1.28, *p*<.001, 95% CI=1.13-1.46), which was similar to full sample. Furthermore, omnibus parametric and nonparametric tests showed the same pattern of results in the European only subsample as with the full sample (eTable 3). That is, significant PGS group differences emerged across most major indices of functioning with the exception of alcohol abuse/dependence, hypertension, and high blood cholesterol. The directions of these differences were also consistent with what was reported in the full sample (see eTable 4).

## Discussion

Results from this investigation yielded the following discoveries. First, individuals at the lowest end of the ADHD PGS distribution (i.e., lowest 20^th^ percentile) had the lowest probabilities of ADHD, particularly in relation to individuals with high PGS (i.e., the highest 20^th^ percentile). Second, individuals with low ADHD PGS had a 17-19% reduction in risk for ADHD relative to the observed 8.3% prevalence rate of ADHD in Add Health, which also tracks with the national prevalence rate among children in the U.S. population (Danielson et al., 2018). Third, individuals with low ADHD PGS demonstrated superior functional outcomes (i.e., higher cognitive performance, greater levels of educational attainment, and lower BMI) relative to individuals representing the rest of the ADHD PGS distribution, including those who were in the medium and high PGS groups. Collectively, these findings indicate that PGS likely capture far more than just the risk and the absence of risk for a psychiatric outcome; where one lies along the PGS distribution may predict diverging functional consequences, for better and for worse.

First, this study provides novel evidence of qualitative differences depending on where one lies on the psychiatric PGS distribution. In the current study, low PGS individuals were not only distinguishable from their high PGS counterparts on ADHD and adult functional outcomes (which was expected), but they were also distinguishable from their medium PGS peers by virtue of their superior functioning on their cognitive abilities, higher educational attainment, and lower BMI. These findings call into question whether it is apt characterize low PGS individuals as having “low genetic risk” (Torkamani et al., 2018) because it does not sufficiently disambiguate those with fewer susceptibility alleles from the vast majority of individuals in the population who carry a modest burden of genetic liability for psychiatric illness (i.e., the medium PGS group). This issue is magnified in recent findings from GWAS, which have shown that psychiatric risks increase across the PGS distribution for ADHD (Demontis et al., 2019) and major depressive disorder (Wray et al., 2018). However, these studies have only shown how the risk for psychiatric illness increases as a function of PGS when the lowest PGS percentile is the reference point. The effect sizes from these comparisons may have been exaggerated because those at the lowest PGS percentiles may be more reflective of “super controls” rather than typical non-clinical controls at the population level (Plomin et al., 2009). Future genetic association studies should consider using more informative (i.e., less biased) reference points when examining the relative psychiatric risk as a function of one’s PGS. For instance, researchers could instead select a point along the PGS distribution that best maps on to the prevalence rate of the outcome in question. The practical utility of this approach was demonstrated in the current investigation.

Another significant discovery from this study is that medium and high PGS individuals also differed in their risk for ADHD as well as on several indices of adult functional outcomes (i.e., higher cognitive abilities and educational attainment, lower rates of illicit drug abuse/dependence, being ever arrested and perceived stress, and lower BMI). This finding may be crucial because it points to the possibility of a PGS ‘clinical threshold,’ as theorized in the classic liability threshold model of psychiatric illness (pp. 36-37, Knopik et al., 2016). The liability threshold model states that while genetic risk is distributed normally in the population, the disorder should only occur once a threshold of liability has been met or exceeded. Individuals who meet or exceed this liability threshold may also carry a different set of causal risk factors that differentiate them from those at the extreme ends of the normal range (Rutter et al., 1996; Rutter and Sroufe, 2000). Thus, it is plausible that observed qualitative differences between medium and high PGS individuals on ADHD risk and adult functional outcomes may reflect not only differences in their genetic risk, but also environmental risks as well. Clearly, future studies aiming to identify a clinical threshold for PGS that draws the line between disorder and non-disorder (or perhaps more apt for those who subscribe to dimensional characterizations of psychiatric illness, distress versus non distress) will be important if PGS is to be useful for clinical practice.

The findings should be interpreted in light of several study limitations. First, the absence of ADHD GWAS featuring well-powered samples of non-European individuals make it challenging to generalize ADHD PGS to populations that are not of predominantly European ancestry, such as Add Health. This limitation was somewhat alleviated by the several methods that were employed to safeguard against the effects of population stratification (i.e., PGS that were standardized by genetic ancestry, PCs covaried in each of the analyses). Additionally, the European-only analyses showed that PGS group differences on the risk for ADHD and adult functional outcomes were almost entirely consistent with what was observed from the full sample. Still, future investigations should anticipate the greater prediction error that comes with using a European-based PGS in mixed samples. Second, the data used in this study were limited to retrospective self-reports, which might be confounded via rater bias or potential issues with validity. In the case of ADHD, participants retrospectively reported on their childhood ADHD symptoms as adults (at Wave III). While adults are fairly reliable reporters of their own ADHD when compared to parents and clinicians (Adler et al., 2006; Kooij et al., 2008), they also tend to underreport the severity of their symptoms (Kooij et al., 2008). Still, having ADHD data from multiple raters would have strengthened this study considerably, especially given that heritability estimates of ADHD differ by rater (Larsson et al., 2014, 2013; Merwood et al., 2013). Finally, this investigation did not assess other disorders that are known to covary with ADHD. More complex phenotypic models that account of the shared genetic underpinnings of co-occurring phenotypes are needed in future studies.

Findings from this study may spur a crucial shift in perspective with regard to the utility of psychiatric PGS in future investigations. Furthermore, PGS may soon play a role in clinical prediction and decision making (Anderson et al., 2019; Bogdan et al., 2018). For instance, it is possible that individuals at high genetic risk may benefit more from an interventions relative to those at moderate and even low genetic risk (Mega et al., 2015; Natarajan Pradeep et al., 2017). At the same time, it may be useful to consider PGS as part of a broad constellation of both risk and protective factors when determining treatment recommendations for ADHD and other psychiatric disorders. Uncertainty in clinical decision making can be reduced with a comprehensive view of patient care that aggregates our increasing knowledge of genetic risks along with crucial clinical information (i.e., biomarkers, environmental factors) pertinent to the individual.

