## Supplemental Materials for "The Positive End of the Polygenic Score Distribution for ADHD: A Low Risk or a Protective Factor?"

eTable 1. Demographic statistics comparing Add Health genetic sample to non-genetic sample
eTable 2. Bivariate correlations
eTable 3. Tests comparing PGS status across functional outcomes for the European subsample only
eTable 4. Pairwise contrasts of PGS groups on dependent variables for the European subsample only (continued on next page)

eTable 1. Demographic statistics comparing Add Health genetic sample to non-genetic sample

| **Variables** | **Genetic sample means (SD) or N's (%)** | **Non-genetic sample means (SD) or N's (%)** | **Test statistic** | ***p*** | **Effect size** |
|---|---|---|---|---|---|
| N | 7088 | 7691 | | | |
| Age | 29.00 (1.74) | 29.18 (1.74) | $t$=6.02 | <.001 | Cohen's $d$=.103 |
| Sex | | | | | |
| Male | 3244 (45.8%) | 3538 (46.0%) | $\chi^2$=5.82 | .016 | Cramer's $V$=.019 |
| Self-reported race | | | $\chi^2$=262.86 | <.001 | Cramer's $V$=.130 |
| Caucasian (including Hispanic) | 4506 (63.6%) | 5343 (66.7%) | | | |
| African American | 1467 (20.7%) | 1627 (20.3%) | | | |
| Native American | 17 (.2%) | 33 (.4%) | | | |
| Asian | 365 (5.1%) | 401 (5.0%) | | | |
| Other | 732 (10.3%) | 286 (3.6%) | | | |
| Missing/Not reported | 1 (<.1%) | 1 (<.1%) | | | |
| ADHD diagnostic criteria | 587 (8.3%) | 576 (7.4%) | $\chi^2$=7.43 | .006 | Cramer's $V$=.024 |
| Cognition | | | | | |
| AHPVT standardized score | 100.57 (14.18) | 98.62 (15.47) | $t$=-7.94 | <.001 | Cohen's $d$=.131 |
| Educational Attainment | | | | | |
| Highest degree attained | 5.62 (2.02) | 5.60 (2.03) | $t$=-.46 | .621 | Cohen's $d$=.010 |
| Mental Health and Behavior | | | | | |
| CES-D depression symptoms | 6.75 (4.59) | 6.78 (4.60) | $t$=-.60 | .550 | Cohen's $d$=.007 |
| DSM-IV alcohol abuse/dependence | 1885 (26.6%) | 2031 (26.4%) | $\chi^2$=17.44 | <.001 | Cramer's $V$=.033 |
| DSM-IV illicit drug abuse/dependence | 542 (7.6%) | 583 (7.6%) | $\chi^2$=1.89 | .169 | Cramer's $V$=.011 |
| Ever arrested | 2055 (29.0%) | 2231 (29.1%) | $\chi^2$=1.15 | .284 | Cramer's $V$=.009 |
| Perceived stress | 4.85 (2.97) | 4.86 (2.98) | $t$=-.27 | .789 | Cohen's $d$=.003 |
| Physical Health | | | | | |
| BMI | 3.32 (1.30) | 3.32 (1.30) | $t$=-1.16 | .248 | Cohen's $d$=.000 |
| Hypertension stage 2 | 905 (12.8%) | 994 (12.9%) | $\chi^2$=.89 | .345 | Cramer's $V$=.008 |
| High blood cholesterol | 556 (7.8%) | 602 (7.8%) | $\chi^2$=2.66 | .264 | Cramer's $V$=.013 |

eTable 2. Bivariate correlations

| | Variable | 1 | 2 | 3 | 4 | 5 | 6 | 7 | 8 | 9 | 10 | 11 | 12 | 13 | 14 |
|---|---|---|---|---|---|---|---|---|---|---|---|---|---|---|---|
| 1 | Biological sex | 1 | | | | | | | | | | | | | |
| 2 | Age | -.050** | 1 | | | | | | | | | | | | |
| 3 | ADHD PGS | .007 | .029* | 1 | | | | | | | | | | | |
| 4 | ADHD diagnostic status | -.097** | .004 | .056** | 1 | | | | | | | | | | |
| 5 | AHPVT standardized score | -.051** | -.052** | -.079** | .008 | 1 | | | | | | | | | |
| 6 | Educational attainment | .126** | -.021 | -.131** | -.071** | .378** | 1 | | | | | | | | |
| 7 | CES-D depression symptoms | .100** | .007 | .054** | .091** | -.138** | -.190** | 1 | | | | | | | |
| 8 | DSM-IV alcohol abuse/dependence | -.125** | -.044** | -.016 | .064** | .189** | .062** | .014 | 1 | | | | | | |
| 9 | DSM-IV illicit drug abuse/dependence | -.050** | -.009 | .036** | .057** | .047** | -.092** | .102** | .228** | 1 | | | | | |
| 10 | Ever arrested | -.269** | .000 | .060** | .101** | -.059** | -.235** | .075** | .205** | .204** | 1 | | | | |
| 11 | Perceived stress | .098** | -.017 | .049** | .078** | -.100** | -.186** | .676** | .027* | .105** | .092** | 1 | | | |
| 12 | BMI classification | .020 | .036** | .074** | .044** | -.078** | -.118** | .064** | -.062** | -.044** | -.034** | .046** | 1 | | |
| 13 | Hypertension stage 2 | -.084** | .012 | .027* | .045** | -.047** | -.057** | .053** | .005 | .020 | .020 | .045** | .218** | 1 | |
| 14 | High blood cholesterol | -.029* | .058** | .011 | .004 | .033** | .037** | .046** | -.005 | -.010 | -.031** | .023* | .110** | .152** | 1 |

eTable 3. Tests comparing PGS status across functional outcomes for the European subsample only (continued on next page)

| | | | | **Test Statistic** | | | **95% Confidence Interval** | |
|---|---|---|---|---|---|---|---|---|
| **Dependent Variable (range)** | **PGS Status** | **Mean** | ***s.e.*** | ***F*, $\chi^2$ *or H*** | ***p**** | ***df*** | **Lower** | **Upper** |
| Cognition | | | | | | | | |
| AHPVT standardized score (50-141) | Low | 106.55 | 6.64 | *F*=16.68 | <.001 | 2 | 105.27 | 107.82 |
| | Medium | 104.22 | .42 | | | | 103.39 | 105.06 |
| | High | 101.67 | .64 | | | | 100.39 | 102.93 |
| Educational Attainment | | | | | | | | |
| Highest degree attained (1-10) | Low | 6.01 | .15 | *H*=79.47 | <.001 | 2 | 5.78 | 6.23 |
| | Medium | 5.65 | .08 | | | | 5.48 | 5.81 |
| | High | 5.17 | .12 | | | | 4.93 | 5.39 |
| Mental Health and Behavior | | | | | | | | |
| CES-D depression symptoms (0-15) | Low | 5.81 | .17 | *H*=9.78 | .008 | 2 | 5.47 | 6.14 |
| | Medium | 6.47 | .11 | | | | 6.25 | 6.68 |
| | High | 6.77 | .21 | | | | 6.36 | 7.18 |
| DSM-IV alcohol abuse/dependence (0-1) | Low | .35 | .02 | $\chi^2$=.40 | .817 | 2 | .32 | .38 |
| | Medium | .33 | .01 | | | | .30 | .35 |
| | High | .29 | .02 | | | | .25 | .33 |
| DSM-IV illicit drug abuse/dependence (0-1) | Low | .08 | .01 | $\chi^2$=23.61 | <.001 | 2 | .06 | .11 |
| | Medium | .09 | .01 | | | | .08 | .10 |
| | High | .12 | .01 | | | | .09 | .14 |
| Ever arrested (0-1) | Low | .26 | .02 | $\chi^2$=21.07 | <.001 | 2 | .22 | .29 |
| | Medium | .28 | .01 | | | | .26 | .31 |
| | High | .33 | .02 | | | | .29 | .38 |
| Perceived Stress (0-16) | Low | 4.34 | .11 | *H*=14.75 | .001 | 2 | 4.11 | 4.56 |
| | Medium | 4.83 | .08 | | | | 4.65 | 4.99 |
| | High | 4.91 | .12 | | | | 4.67 | 5.16 |

eTable 3. Tests comparing PGS status across functional outcomes for the European subsample only (continued from previous page)

| | | | | **Test Statistic** | | | **95% Confidence Interval** | |
|---|---|---|---|---|---|---|---|---|
| **Dependent Variable** | **PGS Status** | **Mean** | ***s.e.*** | ***F, χ2 or H*** | ***p**** | ***df*** | **Lower** | **Upper** |
| Physical Health | | | | | | | | |
| BMI (1-6) | Low | 3.04 | .04 | *H*=21.90 | <.001 | 2 | 2.96 | 3.13 |
| | Medium | 3.27 | .04 | | | | 3.21 | 3.34 |
| | High | 3.40 | .07 | | | | 3.26 | 3.54 |
| Hypertension stage 2 (0-1) | Low | .11 | .01 | $\chi^2$=4.91 | .086 | 2 | .09 | .14 |
| | Medium | .14 | .01 | | | | .12 | .16 |
| | High | .14 | .01 | | | | .11 | .18 |
| High blood cholesterol (0-1) | Low | .09 | .01 | $\chi^2$=1.42 | .492 | 2 | .06 | .11 |
| | Medium | .08 | .01 | | | | .06 | .09 |
| | High | .09 | .01 | | | | .07 | .12 |

*P-values are reported to the thousandsth decimal point due to the Bonferonni correction threshold of $p<.005$.

*Note*. Results show parametric and nonparametric tests of PGS group comparisons on functional outcomes. A one-way ANOVA was performed to examine differences on continuous and normally distributed outcomes. Kruskal-Wallis H tests were used to test for differences on ordinal or continuous outcomes, and chi-square tests were used for binary outcomes. "Low" PGS is the lowest quintile (<20$^{th}$ percentile) of the PGS distribution, "Medium" PGS is the combination of the second, third, and fourth quintiles (21$^{st}$ – 79$^{th}$ percentile) of the PGS distribution, "High" PGS is the fifth quintile (>80$^{th}$ percentile) of the PGS distribution.

eTable 4. Pairwise contrasts of PGS groups on dependent variables for the European subsample only (continued on next page)

| | | Test Statistic | | |
|---|---|---|---|---|
| **PGS Group Contrasts (dummy code)** | **Dependent Variable** | ***T*, $\chi 2$ *or H*** | ***df*** | ***p*** |
| Low (0) versus Medium (1) | Cognition | | | |
| | AHPVT standardized score | *T*=4.22 | 3723 | <.001 |
| | Educational Attainment | | | |
| | Highest degree attained | *H*=27.14 | 1 | <.001 |
| | Mental Health and Behavior | | | |
| | CES-D depression symptoms | *H*=4.81 | 1 | .028 |
| | DSM-IV alcohol abuse/dependence | $\chi^2$=.40 | 1 | .528 |
| | DSM-IV illicit drug abuse/dependence | $\chi^2$=.05 | 1 | .816 |
| | Ever arrested | $\chi^2$=1.38 | 1 | .241 |
| | Perceived stress | *H*=9.18 | 1 | .002 |
| | Physical Health | | | |
| | BMI | *H*=13.57 | 1 | <.001 |
| | Hypertension stage 2 | $\chi^2$=3.06 | 1 | .080 |
| | High blood cholesterol | $\chi^2$=.62 | 1 | .432 |
| Low (0) versus High (2) | Cognition | | | |
| | AHPVT standardized score | *T*=5.61 | 1843 | <.001 |
| | Educational Attainment | | | |
| | Highest degree attained | *H*=77.11 | 1 | <.001 |
| | Mental Health and Behavior | | | |
| | CES-D depression symptoms | *H*=9.59 | 1 | .002 |
| | DSM-IV alcohol abuse/dependence | $\chi^2$=.106 | 1 | .744 |
| | DSM-IV illicit drug abuse/dependence | $\chi^2$=13.95 | 1 | <.001 |
| | Ever arrested | $\chi^2$=17.29 | 1 | <.001 |
| | Perceived stress | *H*=13.85 | 1 | <.001 |

Table 2. Pairwise contrasts of PGS groups on dependent variables for the European subsample only (continued from previous page)

| | | **Test Statistic** | | |
|---|---|---|---|---|
| **PGS Group Contrasts** | **Dependent Variable** | ***F*, $\chi$2 *or H*** | ***df*** | ***p*** |
| Low (0) versus High (2) | Physical Health | | | |
| | BMI | *H*=20.19 | 1 | <.001 |
| | Hypertension stage 2 | $\chi^2$=4.74 | 1 | .030 |
| | High blood cholesterol | $\chi^2$=.05 | 1 | .817 |
| Medium (1) versus High (2) | Cognition | | | |
| | AHPVT standardized score | *T*=2.81 | 3714 | .005 |
| | Educational Attainment | | | |
| | Highest degree attained | *H*=33.72 | 1 | <.001 |
| | Mental Health and Behavior | | | |
| | CES-D depression symptoms | *H*=2.64 | 1 | .104 |
| | DSM-IV alcohol abuse/dependence | $\chi^2$=.05 | 1 | .819 |
| | DSM-IV illicit drug abuse/dependence | $\chi^2$=21.00 | 1 | <.001 |
| | Ever arrested | $\chi^2$=15.89 | 1 | <.001 |
| | Perceived stress | *H*=15.35 | 1 | .125 |
| | Physical Health | | | |
| | BMI | *H*=3.82 | 1 | .051 |
| | Hypertension stage 2 | $\chi^2$=.81 | 1 | .368 |
| | High blood cholesterol | $\chi^2$=1.15 | 1 | .283 |

*P-values are reported to the thousandsth decimal point due to the Bonferonni correction threshold of $p<.005$.
